# Photocontrolled Release of Nitric Oxide for Precise Management of NO Concentration in a Solution

**DOI:** 10.1101/2022.11.01.514656

**Authors:** Egor O. Zhermolenko, Tatyana Yu. Karogodina, Alexey Yu. Vorobev, Mikhail A. Panfilov, Alexander E. Moskalensky

## Abstract

Nitric oxide (NO) is a unique biochemical mediator involved in the regulation of vital processes. Its short half-life and local action impede direct application in medical practice. Light-controllable NO releasers are promising for the development of smart therapies. Here we present novel aza-BODIPY derivative containing *N*-nitroso moiety, which shows efficient and reversible release of NO under red light. Using this molecule, we demonstrate a system for precise management of NO concentration in aqueous solution *in vitro* using the feedback loop with optical control. Our results could serve as the basis for the development of novel therapeutic methods.

## Main

Nitric oxide (NO) is a short-lived, endogenously produced signaling molecule [1]. NO has many physiological roles and is involved in pathological processes as well. Due to short lifetime (∼5s) and limited diffusion range, the potential agents for NO-based therapy should rely on local *in situ* generation. This impedes application of the wide range of NO donors that are currently available for the treatment of cardiovascular diseases, wound healing, immune response to infection, and cancer therapy. Clinical applications are largely limited to the inhaled NO gas [2]. Interestingly, it has already been reported as a potential candidate for use in the treatment of human coronavirus infections, including COVID-19 [3]. However, the action of inhaled NO is limited to lungs, whereas controlling its levels in biological fluids is promising for potential therapeutic applications.

At physiological level of 50-100 nM, NO provides smooth muscle relaxation necessary for normal blood flow [4] and inhibits platelet activation [5,6]. The decreased bioavailability of NO, e.g. due to endothelial dysfunction and/or oxidative stress, is implicated in the development of vascular complications of diabetes mellitus [7,8]. Intermediate concentration of NO promotes growth and proliferation of normal cells and angiogenesis [9]. In contrast, at high concentration both effects are inhibited [10,11]. Conventional NO donors are not suitable for angiogenesis due to their initial burst-release profile. Strategies to slow down the release should be employed, as was successfully demonstrated in [12].

As high concentration of NO provides cytostatic effect, it was hypothesized to be useful in cancer treatment. However, it may have a dual pro- and antitumor action, depending on the local concentration of the molecule [13]. It was shown that NO at the levels ≳300 nM induces the activation of p53, a tumor suppressor protein. An NO-induced apoptosis was observed in cancer cells in several studies [14–16], as well as the inhibition of the immune escape [17]. Intermediate levels of NO can have the opposite, tumor-promoting effect, resulting in the biphasic “double-edged sword” behavior [18]. Therefore, an important problem to be solved for NO-based therapy (including therapeutic angiogenesis and anti-tumor treatment) is the precise control of its concentration.

For controlled delivery of NO, extensive efforts are being made to develop novel NO-releasing biomaterials [19,20] such as liposomes, nanoparticles, macromolecules and dedicated devices [21]. The most promising methods rely on the triggering of NO release using external stimuli, for example, infrared radiation [22], X-ray radiation [23] and ultrasound [24]. In recent years, several molecules capable of NO release under the action of light (“NO photodonors”) were reported. They usually consist of a light-absorbing antenna (a chromophore) attached to an NO-carrying group, such as -NO_2_ in dimethylnitrobenzenes (DNB) [25–27] or *N*-nitroso-group [28,29]. The photodonors working in red or infrared region are of particular interest because such a light penetrates deep into biological tissues. Recently, NO-photodonors based on aza-BODIPY core conjugated with N-NO fragment have been reported [30,31]. Aza-BODIPY dyes have superior spectral performances, such as red-shifted absorption spectra and high molar extinction coefficients, and are considered to be extremely attractive organic small-molecule materials for various bioapplications [31,32].

In this paper, we present novel relatively simple NO photodonor based on aza-BODIPY core bearing two N-NO units. We show that it can reversibly release NO under near-infrared light, providing stationary concentration related with the light intensity. Based on this ability, we implemented an approach for precisely controlled photorelease of NO, providing the desired continuous concentration in an aqueous solution. We use the feedback loop consisting of ISO-NOP sensor for real-time determination of NO concentration and a photodonor which is illuminated by laser beam. The compartment containing photodonor is separated from the sensor compartment by gas-permeable membrane, resulting in clean NO generation without any additional chemicals. Our approach demonstrate successful controllable generation of NO and active maintenance of its concentration, which is a crucial step towards the NO-based smart therapies.

## Results

### Synthesis and NO Photorelease Studies

Synthesis of the **AzaB-NO** was provided according to the Scheme 1. We used the approach described in [30] to obtain the nitro derivative **1**. Next, in contrast to above the mentioned paper, we put **1** into homo-condensation with ammonium acetate yielding of aza-dipyrromethene **2**. Its complexation with boron trifluoride and nitrosation leads to the target product **AzaB-NO**. It has symmetrical structure with two –NO tails and therefore is potantionally capable of releasing two NO molecules under light. More detailed description of synthetic procedures and characterization of the molecule can be found in the Supplementary materials.

**Scheme 1.**
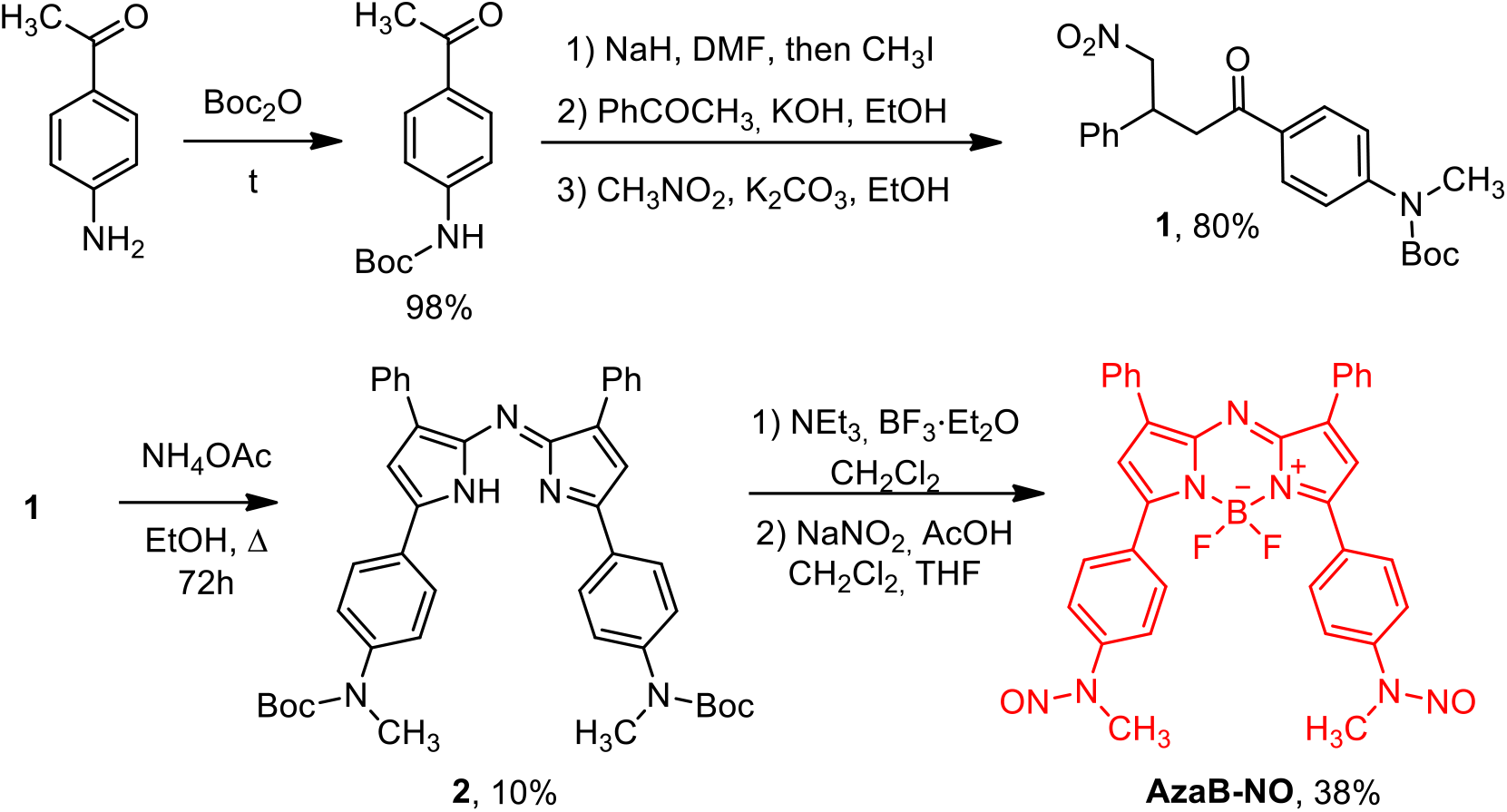
Synthesis of **AzaB-NO**.

The absorption spectra and its changes upon irradiation are shown on Figure 1A. Initially, **AzaB-NO** has strong absorption peak at 672 nm in EtOH (shown by red line). During illumination with 660 nm light, the transformation of **AzaB-NO** to another form occurs as the new absorption peak appears at a wavelength of 736 nm, while an isobestic point is observed at 700 nm. This fact is consistent with the literature data for similar compounds, where this transformation was correlated with the release of NO [30,31]. No significant fluorescence was observed for **AzaB-NO**, apparently due to intramolecular photoinitiated electron transfer from N-NO unit to aza-BODIPY core.

**Figure 1.**
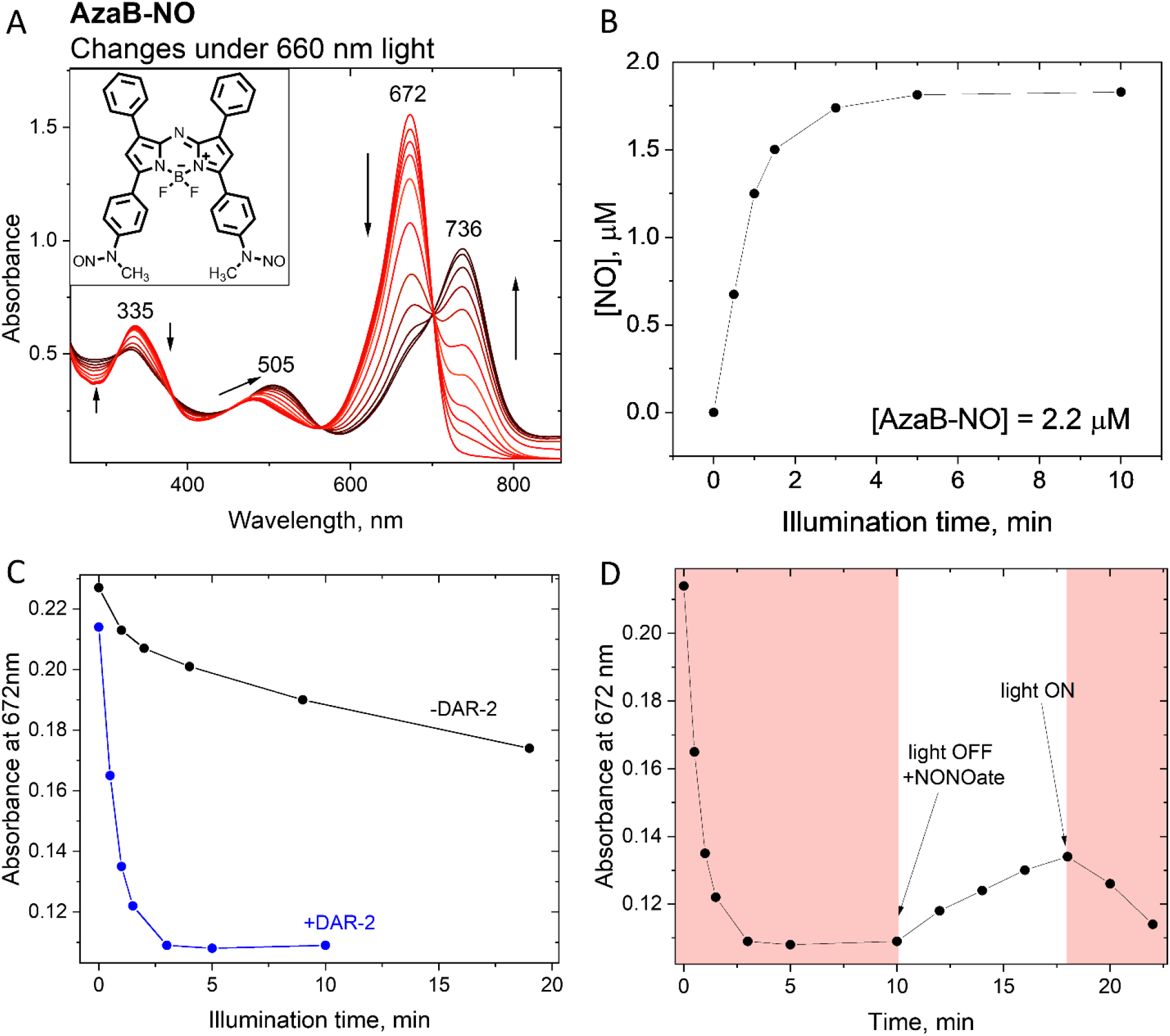
A. The structure of **AzaB-NO** photodonor and changes of its absorption spectrum during illumination with 660 nm LEDs. B. Release of NO measured during illumination using DAR-2 fluorescence probe. C. Comparison of the time course of spectral change (decrease of the 672 nm absorption peak) with and without DAR-2. D. Demonstration of the reversibility of NO photorelease using NONOate.

The release of NO for **AzaB-NO** was estimated using the DAR-2 fluorescent probe [33,34]. 2-fold amount of DAR-2 was added to the solution of **AzaB-NO** in EtOH, and fluorescence (λ_exc_=540 nm) was measured after each minute of photolysis. The huge, 20-fold increase of fluorescence was observed (whereas there is no intrinsic fluorescence of the photodonor at this excitation wavelength). The fluorescence was recalculated to the NO concentration using the approach described in Supplementary. The results are shown in Figure 1B. The amount of detected NO corresponds to ∼80% of the initial concentration of **AzaB-NO** in a sample, whereas theoretical maximum is 200% (two NO molecules can be released by each **AzaB-NO**). However, it is natural to assume that not every molecule is captured by DAR-2 probe because other concurrent reactions can occur.

Interestingly, the changes in absorption spectrum occur much faster with the presence of DAR-2 probe (Figure 1C). Obviously, this is due to the irreversible removal of NO from the system. This led us to the conclusion that NO could attach itself back to the **AzaB-NO** effectively slowing down the release. Indeed, the *N*-nitrosation was shown for similar compounds [35]. To test directly the possibility of the reverse reaction, we added the conventional NO donor (NONOate) to the reaction mixture and observed the backward change of absorption spectrum. Further illumination led to the same (“forward”) effect as before (Figure 1D). The reversibility is also confirmed by HPLC experiments (see supplementary materials).

**Scheme 2.**
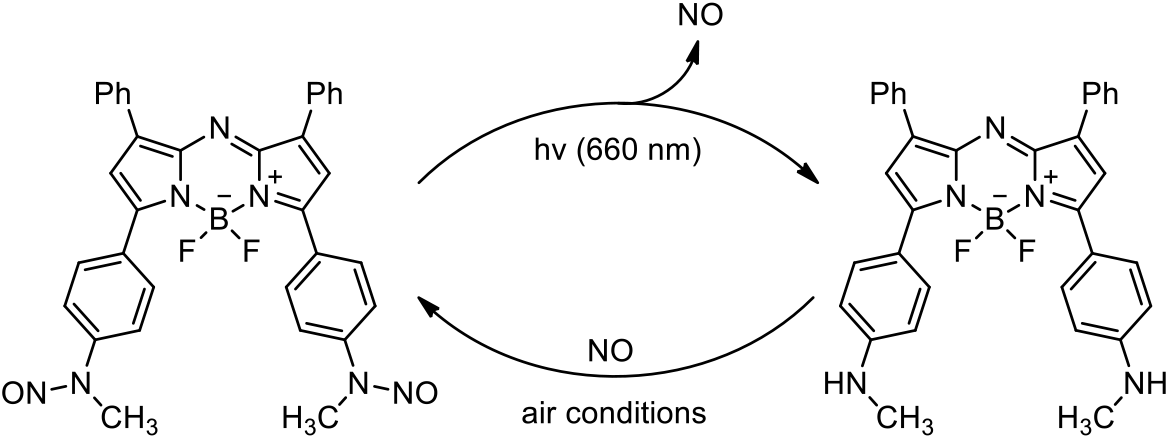
The scheme of photoinduced reactions for **AzaB-NO**.

Therefore, the photolysis of **AzaB-NO** is reversible and occurs according to Scheme 2. This property is interesting and allows one to use this molecule not only as NO source but also as buffering system capable of capturing NO in the dark for subsequent release under light.

### Feedback system

A challenging problem to be solved prior to biological application of the NO photodonors based on aza-BODIPY core is poor solubility and precipitation in aqueous solution. This problem is often ignored or solved partly by using emulsifiers or encapsulating the dye into micelles [31,36]. In contrast, we decided to use the gaseous nature of NO, i.e., its ability to penetrate through gas-permeable membranes. In this concept the solution of the NO photodonor in an organic solvent is placed on one side of the membrane, whereas aqueous solution is on the other side. In such a system one can use huge concentration of the photodonor, which provides wide dynamic range of NO concentration. Another advantage is that no molecules except NO could transfer through the membrane (in an ideal case). This is especially important for photodonors which are *N*-nitrosamines: at least some compounds of this class are known to be cancerogenic [37]. As a solvent, we chose DMSO because it is biologically inert and has slow evaporation rate, which minimizes the amount of solvent gaseous molecules able to pass through the membrane.

Figure 2 shows the scheme of experimental setup for the photoinduced generation of NO controlled with feedback loop. A beam of 660 nm, 50 mW CW laser (CrystaLaser DL660-050) goes through the 1 mM solution of **AzaB-NO** in DMSO. The mean laser power is modulated by PWM output of Arduino microcontroller connected to TTL input of the laser controller. ISO-NOP sensor is used to measure the NO concentration. Its diameter (2 mm) coincides with the laser beam size, that is why we use it without focusing. The sensor is placed into the Faraday cage to avoid the influence of external electric fields. Right part of the Figure 2 shows the close-up view of the sensor tip and the area below. The sensor is submerged in a capillary filled with the acidic aqueous solution (0.1 M of H_2_SO_4_ in H_2_O). The acidic conditions are needed to increase the lifetime of NO. The tip of the sensor rests on the tensioned layer of Parafilm® which plays a role of gas-permeable membrane, separating the content of the capillary and the organic phase below. Laser beam directly below the membrane induces the release of free NO which goes to the sensor, resulting in the increase of current. The current is converted to voltage by the WPI TBR 1025 analyzer, and this voltage is in turn measured by ADC and read by Arduino microcontroller. The microcontroller transmits data to PC and receives back commands to manage the laser power. More detailed technical description of the setup can be found in Supplementary materials. Finally, to avoid confusion, we note that the sensor has its own NO-selective membrane, which is not compatible with organic solvents. This is another reason why we have to use another membrane – to protect the sensor from DMSO. In tests prior to experiments, we did not observe any influence of DMSO placed below the Parafilm® layer on the sensor readings. The laser beam directed at the sensor tip also did not alter the current.

**Figure 2.**
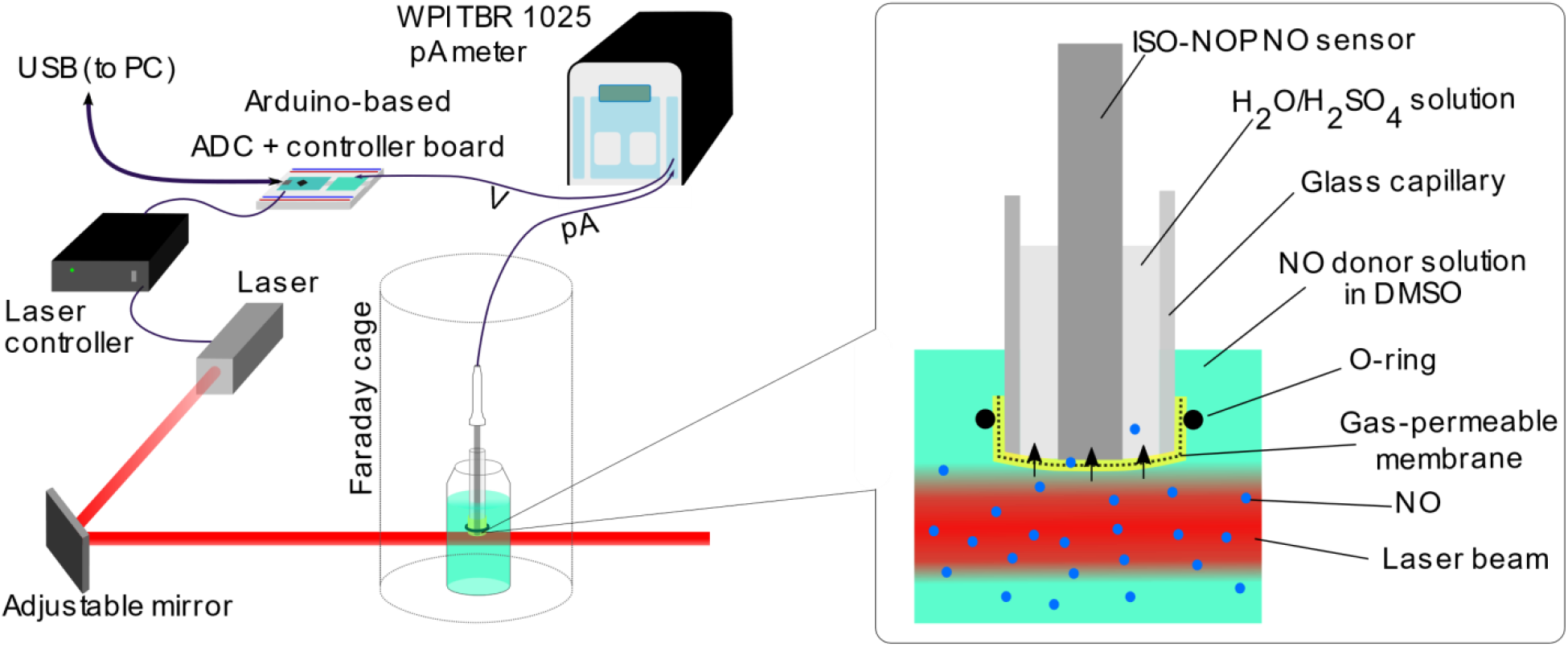
An experimental setup for the photoinduced generation of NO controlled with feedback loop. Left: general scheme; Right: close-up view of the sensor which is separated from the organic phase with NO photodonor by gas-permeable membrane.

Figure 3A shows the sensor readings during multiple cycles of laser irradiation (30 s on / 30 s off). The baseline current is usually stable with a precision of ±1 pA. In contrast, the current increases rapidly by tens of pA as the laser turns on and then decreases back as the laser turns off. This is the result of NO photorelease and subsequent diffusion through the membrane. To prove this point, we performed the control experiment with tetraphenyl-aza-BODIPY (TP-aza-BODIPY) with matched optical density and observed small but visible response (Figure 3B). It can be explained by the fact that TP-aza-BODIPY effectively absorbs 660 nm light and increases the temperature of the solution, which might influence the sensor reading. Although this influence is small, we performed additional experiments with temperature sensor as well. Figure 3C shows parallel measurements of NO concentration and temperature for a solution of **AzaB-NO** during single 1 min illumination session. While temperature response is monotonic and the onset is delayed, the ISO-NOP current shows several phases, designated as 1-5 (which can also be seen on the Figure 3A except 3, due to shorter illumination time). Immediate increase of current (phase 1) is followed by mild decrease (phase 2) and the equilibration (phase 3). Surprisingly, after turning off the laser, the current drops below the initial baseline value (phase 4) and then slowly returns back (phase 5).

**Figure 3.**
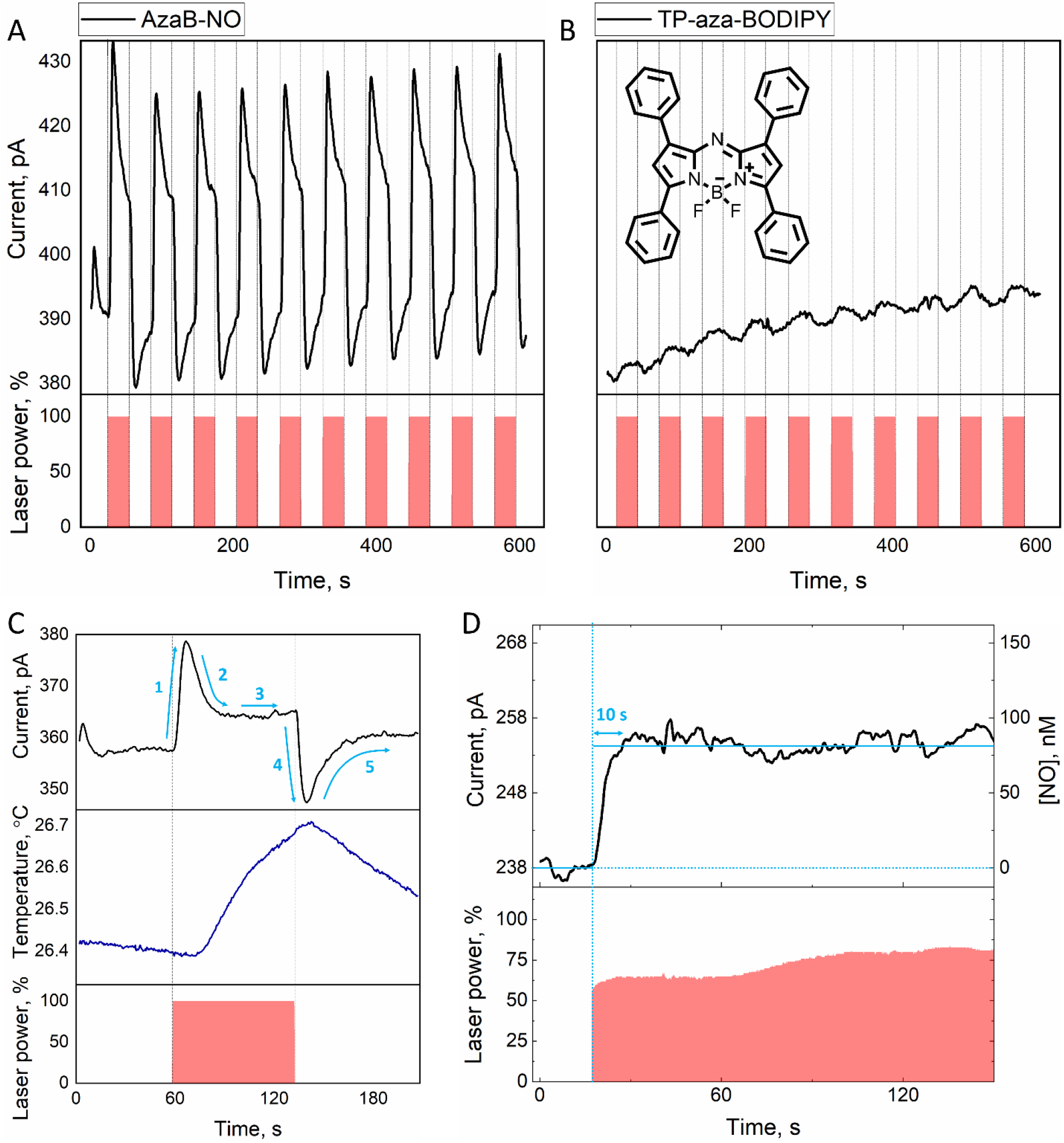
A: Sensor readings during multiple cycles of laser irradiation (30 s on / 30 s off). B: The same for the control compound without NO – tetraphenyl-aza-BODIPY (TP-aza-BODIPY). C: NO sensor readings and temperature before, during and after 60 s irradiation session. D: An illustration of the PID controller in action: rapid increase of NO concentration to the desired level (80 nM) and sustaining by tuning the laser power.

Obviously, during sharp initial burst of NO its concentration can significantly exceed the stationary one, which could result in undesirable (e.g., opposite) effects for biological systems. In the following experiment, we used the power of a feedback loop – the ability to control the laser power dynamically to sustain the concentration at desired level. Specifically, we implemented the PID controller in a dedicated script on a PC. Details of the algorithm can be found in Supplementary materials. Figure 3D shows an illustration of the PID controller in action: rapid increase of NO concentration to the desired level (80 nM) and sustaining by tuning the laser power.

## Discussion

The non-monotonic response of the sensor to the laser irradiation (Figure 3C) should be explained. First, let us discuss the effect which the laser beam has on the solution of **AzaB-NO**. Assuming the linear response, the rate constant of NO photorelease is *k*_1_ = *α* · *I*_0_, where *α* is the proportionality constant and *I*_0_ is the intensity of light at the point of consideration. The simplest kinetic scheme accounting for the reverse reaction is:

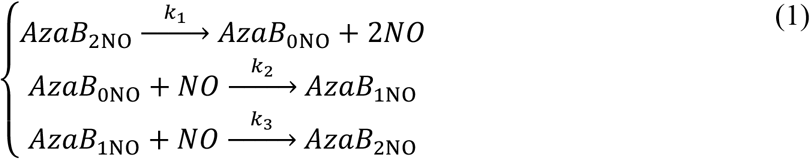

where *AzaB*_2NO_ denotes the concentration of **AzaB-NO**, and the subscript indicates the number of *N*-nitroso groups left on the molecule and *k*_2_ and *k*_3_ are rate constants of reverse reactions. The scheme (1) gives the following expression for the stationary concentration of NO (see supplementary materials for the derivation and discussion of limiting cases):

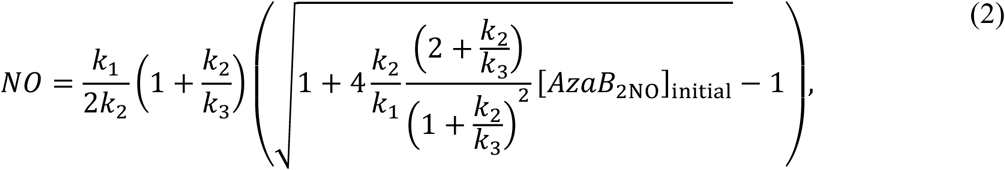

where [*AzaB*_2NO_]_initial_ is the initial concentration of **AzaB-NO**. Importantly, [*NO*] could be much smaller than [*AzaB*_2NO_]_initial_, meaning that only a small fraction of NO exists in a free form, while other molecules are being constantly captured due to the buffering capacity of the photodonor. Eq. (2) gives the correct estimation of initial NO concentration after turning on the laser, but is does not account for the NO degradation and diffusion which are slow but important processes. The initial increase of the sensor readings (phase 1) is than related to the diffusion of NO molecules through the gas-permeable membrane. Maximal response which we observed in our experiments corresponds to ∼300 nM of NO, but it can be further increased by the increase of [*D*_*NO*_]_0_ and/or intensity of the laser beam (for instance, introduction of the focusing optics). The subsequent decrease (phase 2) reflects the change of [*NO*] in the illuminated area, which can be related with different processes: NO reaction with oxygen, diffusion to the unilluminated area, changes of [*D*_*NO*_]/[*D*] balance due to diffusion of aza-BODIPY, and change of the intensity of laser beam due to the change of absorption prior to the area of interest. All these factors could influence the stationary concentration of NO. As this stationary [*NO*] is achieved, it is visible as the stabilization of the sensor current (phase 3). When the laser is turned off, all the free molecules of NO are captured in the illumination area due to reverse reaction. It results in the opposite diffusion flow (from the sensor compartment to the organic phase), which is visible as phase 4. Interestingly, the current drops below the initial value. However, this is not surprising because the results of Figure 3A are actually not from the first experiment, meaning that some non-zero NO concentration existed near the sensor before turning on the laser. When the **AzaB-NO** becomes saturated again, the current returns to the initial value (but is now slightly larger due to residual NO molecules).

Such complex multi-stage time-dependence of the NO concentration in the aqueous solution (i.e., the upper compartment) is unacceptable for biological application. In contrast, the use of feedback loop and PID controller allows one to obtain much predictable behavior, with the concentration of NO rapidly reaching the desired level and then being maintained constant (Figure 3D). First, it should be noted that we used non-classical PID controller but the “proportional-on-measurement” (POM) version instead [38]. Unlike classical PID controller, the POM uses the term *y*(*t*) − *y*(*t*_0_) in place of proportionality term *y*(*t*) − *y*_set_ (where *y* is the sensor readings, *t*_0_ is initial time, *t* is current time and *y*_set_ is the desired value of *y*). In brief, this approach allows one to eliminate the “overshoot”, which was the biggest problem in our experiments. The feedback algorithm contains three parameters (gains for proportional, differential and integral terms) which were tuned manually to provide fast and precise response. The overall system of hardware and software which we developed provides measurement of NO concentration with the frequency of 5 Hz and user interface (on PC) with the ability to plot and control [*NO*] in real time. This system can be easily automated to shape virtually any time-dependency of [*NO*] for long-term studies, for instance, for experiments with cell cultures.

One of the main features of our experiments is that the compartment containing photodonor is separated from the sensor compartment by gas-permeable membrane, resulting in clean generation of NO generation without any additional chemicals. Together with the ability to precisely control NO concentration, this is a crucial step towards the NO-based smart therapies because otherwise the photodonor and its residuals could exert toxic or even cancerogenic effects [37]. Moreover, highly reactive singlet oxygen, which is unavoidably formed during photoexcitation of dyes, cannot pass the membrane due to its low lifetime. The use of special compartment with organic solvent makes it possible to use high concentration of the photodonor, impossible in biological medium, especially in the case of poorly soluble aza-BODIPY dyes. This fundamentally increases the light absorption and the overall efficacy of the system.

The presented concept could be readily tested as a modality for external NO delivery, e.g. transdermal or wound-healing means [39]. Next steps might be the use capsules which can be placed internally or various particles/biomaterials analogous to reported earlier [40]. The presented photodonor is activated by red/near-infrared light with the superior penetration into biological tissues. However, this approach requires additional testing to prevent leaking of the organic phase in the internal environment of a body.

Last but not least, in this paper we report the use of electrochemical sensor which is designed for aqueous solution for the quantification of NO release in an organic solvent. Although it seems to be solely technical advancement, we believe that the reported approach would increase the reliability of the existing data. Usually, the use of ISO-NOP for compounds which are water-insoluble implies the addition of a small percentage of organic solvent, which compromises the lifetime and accuracy of the sensor, and still does not allow to use high concentration of the compound. Although gas-permeable membrane is a trivial solution and was used in several papers for NO quantification [41–44], previously there were no reports on the use of cheap and widespread Parafilm® as a protective membrane for NO sensors.

## Methods

### Photochemical characterization of AzaB-NO

Absorption spectra were measured with Shimadzu UV-1900 spectrophotometer, whereas fluorescence emission spectra were measured with Shimadzu RF-6000 fluorometer.

Photolysis was done in a quartz cuvette using six high-power 660 nm LEDs. LEDs were fixed directly near the cuvette and were fed with the current of 0.25 A.

Photoinduced generation of nitric oxide was evaluated using fluorescent probe DAR-2.

### Electrochemical measurements

For electrochemical measurements we used the WPI TBR-1025 analyzer with ISO-NOP nitric oxide sensor. The sensor was calibrated prior to experiments as described in supplementary. We also performed an additional calibration procedure for sensor protected by the Parafilm® membrane and found that the sensitivity is approximately 5-time lower but still good. In a separate experiments, we found that such a protected sensor does not react to 100% DMSO.

### Electronics

Output of WPI TBR-1025 free radical analyzer is the analogous voltage. An ADS1115 16-bit ADC module was used to convert it into the digital value. The ADC module was connected to Arduino Nano board using standard jumper wires and solderless breadboard. Arduino microcontroller was set up to establish the serial-port connection with PC via the USB cable and to send the digitized sensor readings upon request. Additionally, one of pulse-width modulation (PWM) pins of Arduino microcontroller was connected to the TTL input of the laser controller. The output of this pin was controlled by special command arriving from PC. The firmware for the microcontroller and the overall hardware connections are listed in the supplementary materials.

### Feedback algorithm & software

The software to read data and control the setup was developed as Python script with graphical interface based on `tkinter` and `matplotlib` libraries. The script implements connection with Arduino microcontroller via serial port, requests of data (sending `m` and `t` letters for NO sensor and temperature sensor, respectively), continuous writing of data to file, interactive plotting, switching the laser on/off with user-defined power, and automatic control of NO concentration using PID algorithm with user-defined parameters (including the desired level of concentration). Specifically, we used the `simple_pid` library for Python [45]. The reading of data from Arduino microcontroller and processing of graphics (i.e., data plotting) were implemented in parallel threads to speed up the measurement frequency. The Python script is available upon request.

## Acknowledgements

The study was supported by the Russian Science Foundation (grant #18-15-00049), https://rscf.ru/en/enprjcard?rid=21-15-28006.

## Author contributions

E. Zh.:development of the feedback system, experiments and data collection; T. K: photochemical characterization of the compounds; A.V.: design and synthesis of the compound; A.M. conceptualization and study design, preparation of the manuscript. All authors contributed to the data analysis and provided feedback on the manuscript.

## Competing interests

The authors declare no competing interests.

## Notes

### Competing Interest Statement

The authors have declared no competing interest.

