## Supplementary information for "Photocontrolled Release of Nitric Oxide for Precise Management of NO Concentration in a Solution"

#### Synthesis.

##### General information.

Starting materials unless otherwise noted were obtained from commercial supplies and used without purification.

NMR spectra were recorded on Bruker Avance-300 (300.13 MHz for $^{1}H$) and Avance-400 (400.13 MHz for $^{1}H$ and 100.62 MHz for $^{13}C$) spectrometers, using the residual proton and carbon signals of or $CDCl_3$ ($\delta_H$ 7.24 ppm; $\delta_C$ 77.16 ppm) as internal standards. $^{13}C$ NMR spectra were registered with C-H spin decoupling. Masses of molecular ions were determined by HRMS on a DFS Thermo scientific instrument (EI, 70 eV). UV-VIS spectra was obtained with Shimadzu UV-1900 spectrophotometer. Fluorescence spectrum was recorded with Shimadzu RF-6000 spectrofluorometer.

Reactions were controlled with thin-layer chromatography (TLC). The TLC was carried out on Sorbfil silica plates (UV 254) with further UV light visualization. Flash column chromatography was performed on silica gel (Macherey Nagel, pore size 60 Å, 230–400 mesh).

##### ***tert*-Butyl (4-Acetylphenyl)carbamate.**

*tert*-Butyl (4-Acetylphenyl)carbamate was prepared according to procedure in [1] in 98% yield.

$^{1}H$ NMR (300 MHz, $CDCl_3$, δ): 1.50 (s, 9H); 1.54 (s, 3H); 6.81 (br s, 1H); 7.45 (d, $J$ = 8.6 Hz, 2H); 7.88 (d, $J$ = 8.6 Hz, 2H) [1].

##### ***tert*-Butyl Methyl (4-(4-Nitro-3-phenylbutanoyl)phenyl)carbamate (1).**

*tert*-Butyl Methyl (4-(4-Nitro-3-phenylbutanoyl)phenyl)carbamate was prepared according to procedure in [1] in 80% yield for 3 steps.

$^{1}H$ NMR (300 MHz, $CDCl_3$, δ) 1.46 (s, 9H), 3.28 (s, 3H), 3.40 (dd, J = 7.0, 5.0 Hz, 2H), 4.20 (p, J = 7.1 Hz, 1H), 4.67 (dd, J = 12.5, 7.9 Hz, 1H), 4.81 (dd, J = 12.5, 6.6 Hz, 1H), 7.29 (m, 7H), 7.86 (m, 2H) [1].

##### ***tert*-Butyl (Z)-(4-(2-((5-(4-((tert-butoxycarbonyl)(methyl)amino)phenyl)-3-phenyl-1H-pyrrol-2-yl)imino)-3-phenyl-2H-pyrrol-5-yl)phenyl)(methyl)carbamate (2).**

Mixture of **1** (0.6g, 1.5 mmol**)** and $NH_4OAc$ (1.74g, 22.6 mmol) in EtOH (35ml) was refluxed for 72 hours. The reaction mixture was cooled in a freezer and filtered. Precipitate was washed with 50 ml of cold ethanol. Dark blue solid, yield 53 mg (10%).

$^{1}H$ NMR (300 MHz, $CDCl_3$, δ): 1.50 (s, 18H); 3.34 (s, 6H); 7.15 (s, 2H); 7.31-7.36 (m, 2H); 7.38-7.44 (m, 10H); 7.84-7.88 (m, 4H); 8.01-8.05 (m, 4H); 12.5 (very broad singlet, 1H).

$^{13}C$ NMR (100 MHz, $CDCl_3$, δ): 28.5; 37.2; 81.1; 115.0; 125.5; 126.9; 128.1; 128.4; 128.8; 129.2; 133.9; 142.6; 145.5; 149.8; 154.5.

HRMS: 707.3475 - found, 707.3472 – calculated for $C_{44}H_{45}N_5O_4$.

**5,5-Difluoro-3,7-bis(4-(methyl(nitroso)amino)phenyl)-1,9-diphenyl-5H-dipyrrolo[1,2-c:2',1'-f][1,3,5,2]triazaborinin-4-ium-5-uide (AzaB-NO).**

Compound **2** (80mg, 0.1mmol) was dissolved in 5 of DCM and flushed with argon. DIPEA (0.5 ml) were added and the reaction was stirred for 15 minutes. $BF_3{\cdot}OEt_2$ (0.5 ml) was added dropwise and the reaction was stirred overnight at room temperature. Upon completion, reaction mixture was diluted with EtOAc and subsequently washed with water, 5% $NaHCO_3$ solution and brine. Organic phase was dried over sodium sulfate, evaporated and passed through silica gel column with $CHCl_3$ as eluent. Obtained intermediate (0.19g, 0.03 mmol) was transferred to 25 ml round bottom flask with $NaNO_2$ (0.035 g, 0.05 mmol), DCM (4ml), THF (8 ml) and AcOH (4 ml). Reaction mixture was stirred for 2 hours at room temperature and became dark green. Obtained solution was diluted with DCM and neutralized with 5% $NaHCO_3$ solution and washed with water. Organic phase was dried over sodium sulfate, concentrated and purified by silica gel column chromatography with $CHCl_3$ as eluent. Dark green solid, yield 23 mg (38% over 2 steps).

$^1$H NMR (300 MHz, $CDCl_3$, δ) 3.45 (s, 6H), 7.08 (s, 2H), 7.45 (m, 6H), 7.69 (d, J = 8.5 Hz, 4H), 8.11 (m, 8H).

$^{19}$F NMR (282 MHz, $CDCl_3$, δ) -131.15 (m, 2F).

$^{13}$C NMR (100 MHz, $CDCl_3$) δ 30.3, 118.1, 118.9, 128.6, 129.3, 129.6, 129.8, 130.9, 132.0, 143.8, 144.3, 145.8, 157.9.

HRMS: 613.2213 - found, 613.2209 – calculated for $C_{34}H_{26}{}^{11}BF_2N_7O_2$.

[1] Zhou, E.Y.; Knox, H.J.; Reinhardt, C.J.; Partipilo, G.; Nilges, M.J.; Chan, J. Near-Infrared Photoactivatable Nitric Oxide Donors with Integrated Photoacoustic Monitoring. *J. Am. Chem. Soc.* **2018**, *140*, 11686–11697, doi:10.1021/jacs.8b05514.

### NMR Spectra of obtained compounds

$^1$H NMR spectrum of ***tert*-Butyl (4-Acetylphenyl)carbamate:**

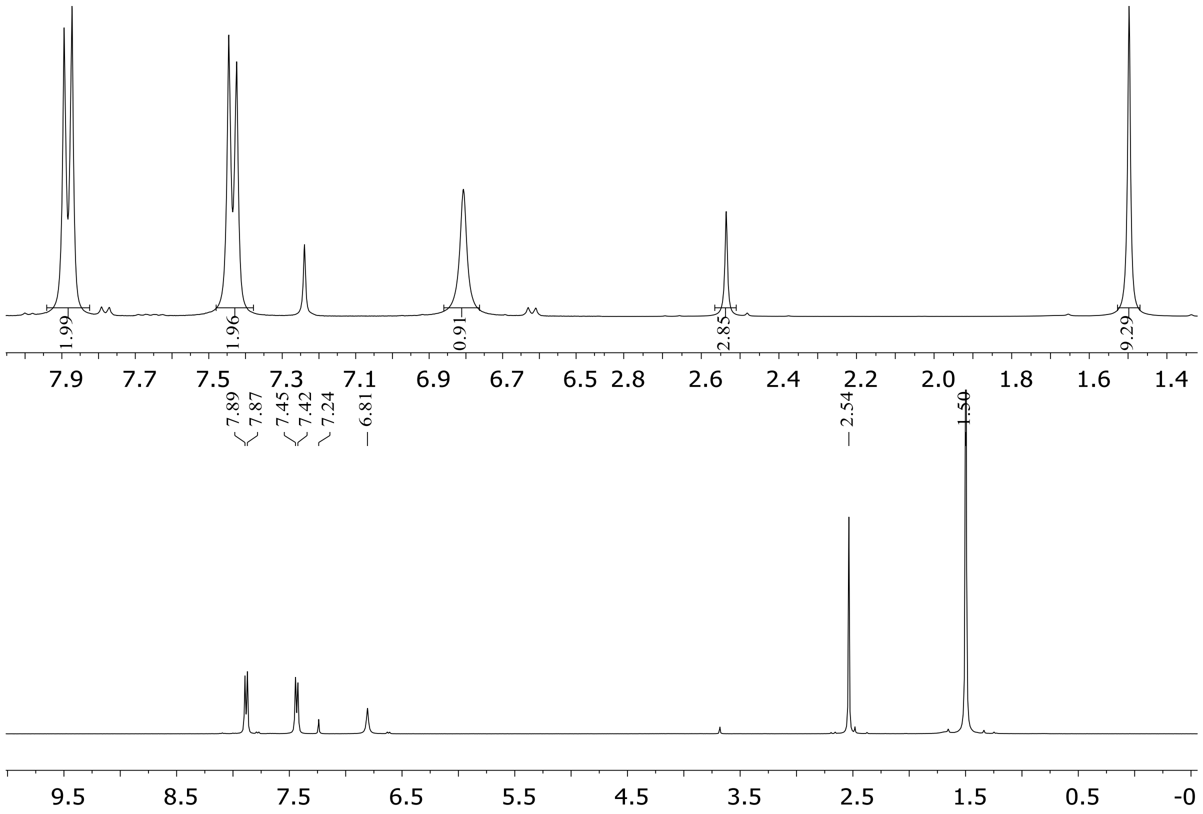

[1]H NMR spectrum of ***tert*-Butyl Methyl (4-(4-Nitro-3-phenylbutanoyl)phenyl)carbamate (1):**

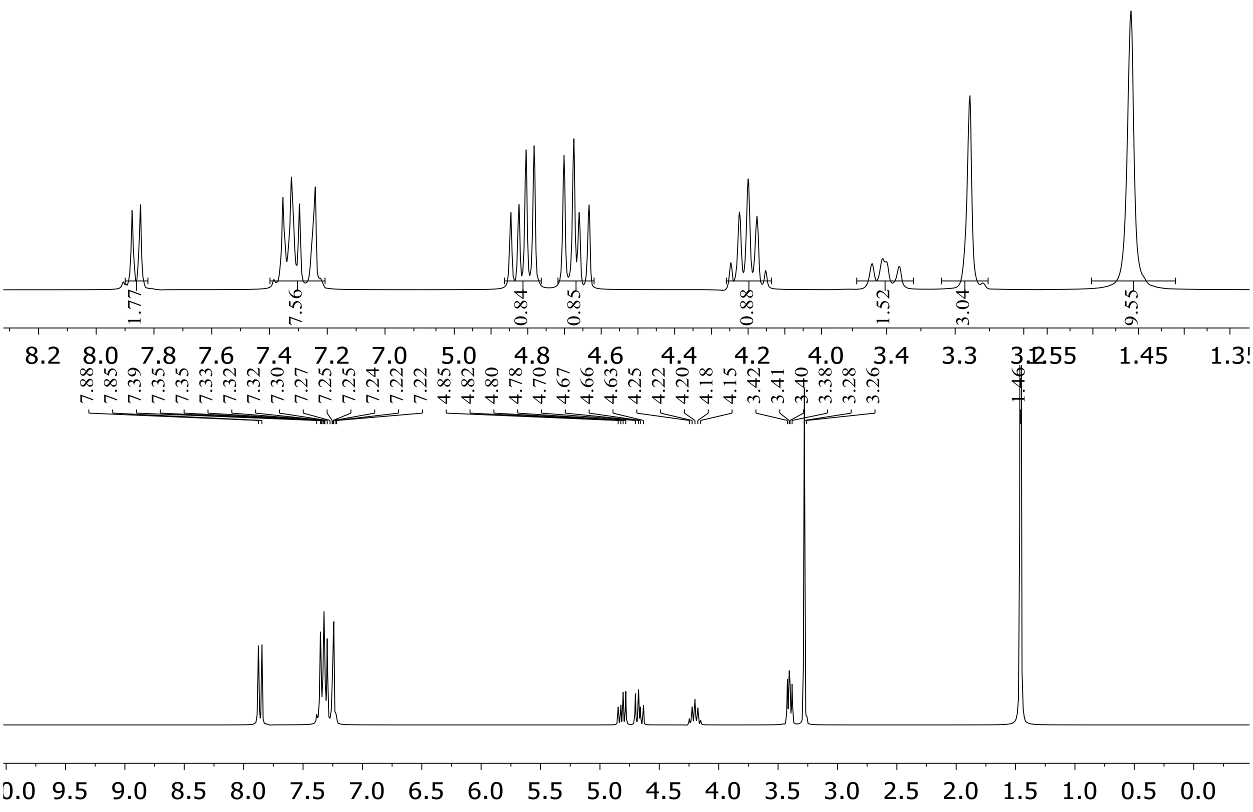

[1]H NMR spectrum of ***tert*-Butyl (*Z*)-(4-(2-((5-(4-((*tert*-butoxycarbonyl)(methyl)amino)phenyl)-3-phenyl-1*H*-pyrrol-2-yl)imino)-3-phenyl-2H-pyrrol-5-yl)phenyl)(methyl)carbamate (2):**

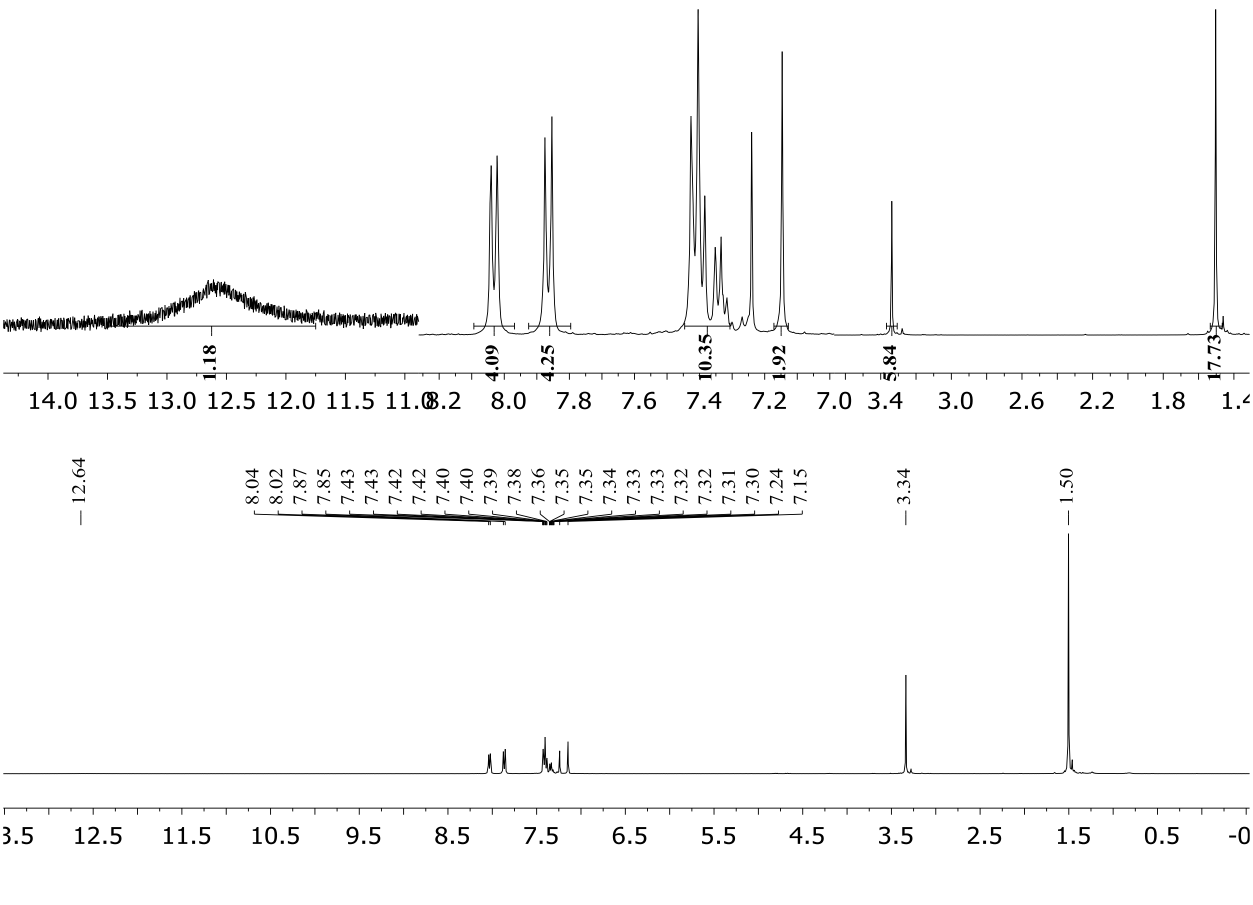

$^{13}C$ NMR spectrum of ***tert*-Butyl (*Z*)-(4-(2-((5-(4-((*tert*-butoxycarbonyl)(methyl)amino)phenyl)-3-phenyl-1*H*-pyrrol-2-yl)imino)-3-phenyl-2H-pyrrol-5-yl)phenyl)(methyl)carbamate (2):**

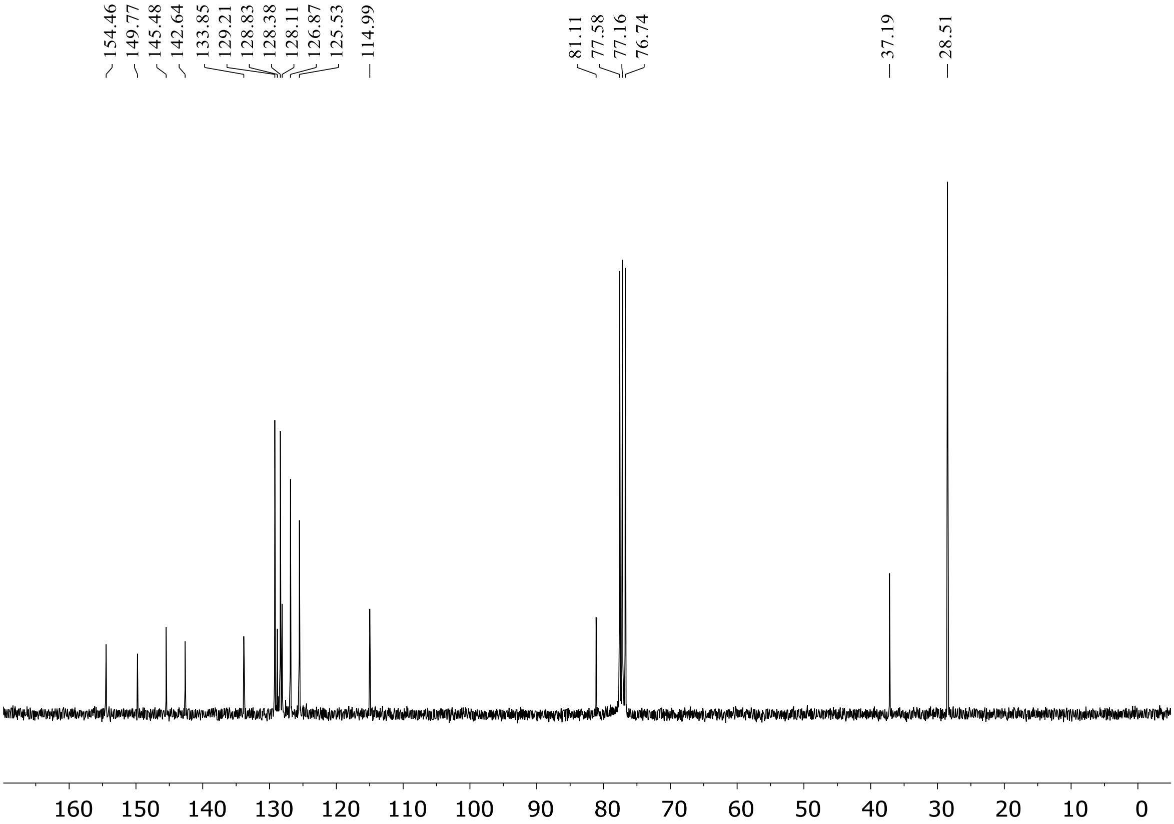

[1]H NMR spectrum of **AzaB-NO:**

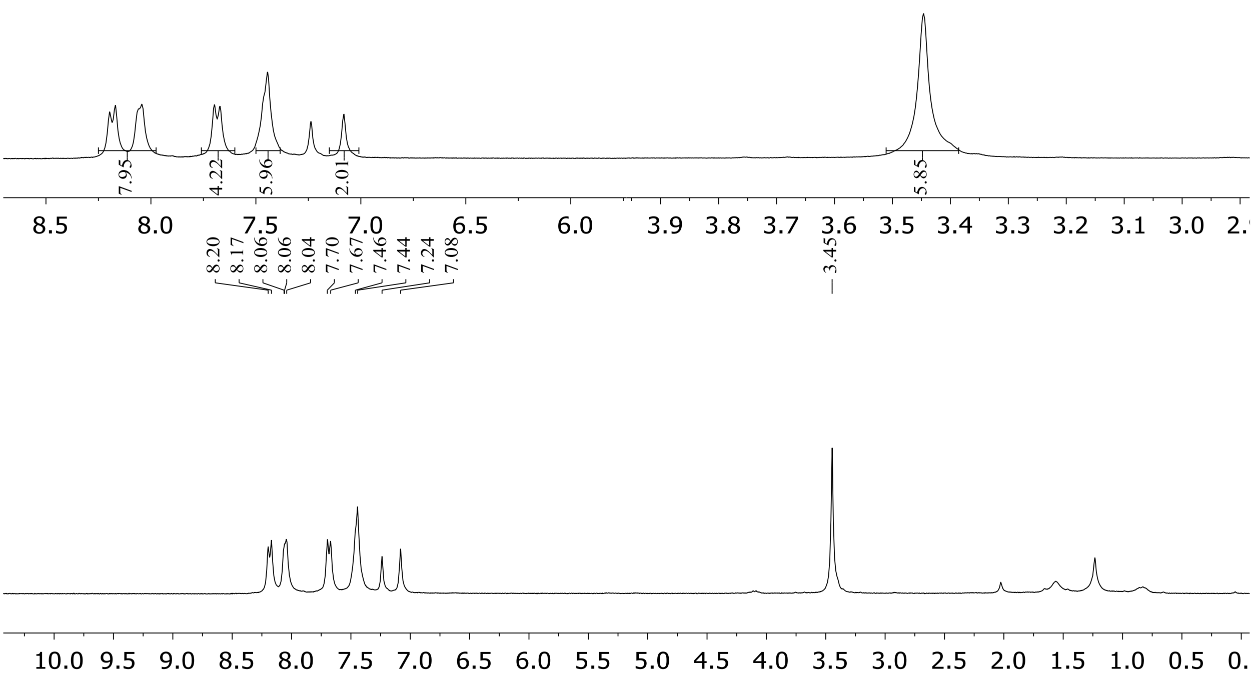

[19]F spectra of **AzaB-NO**:

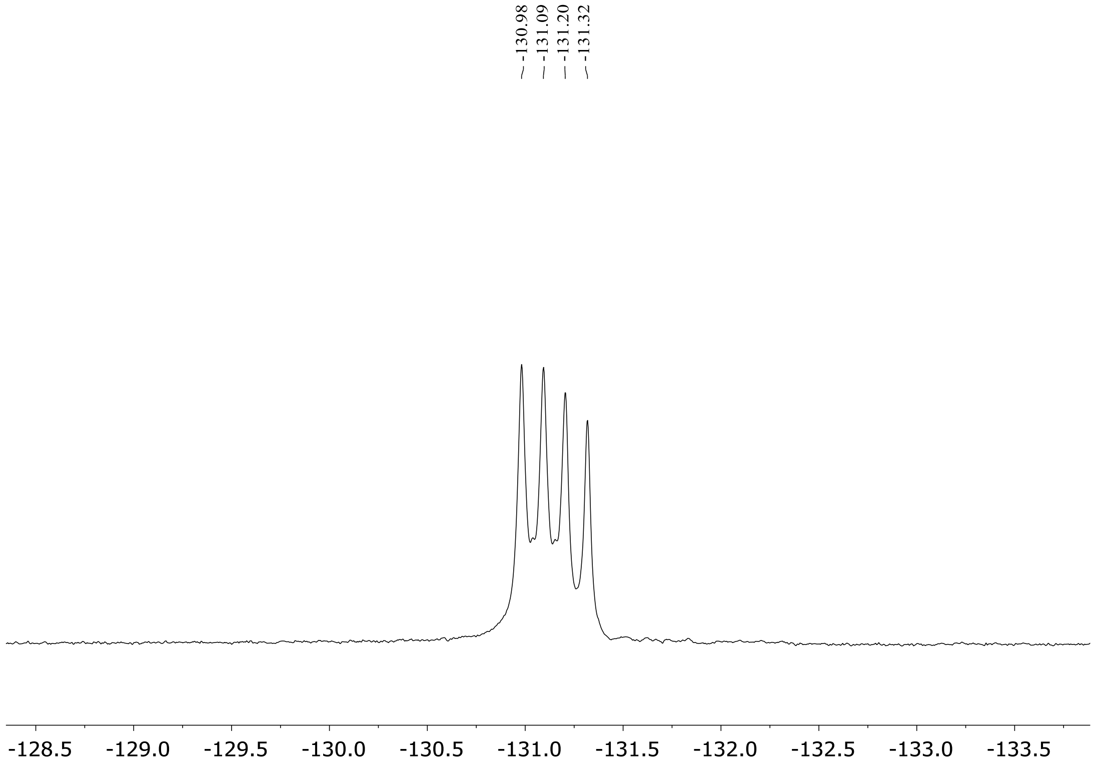

$^{13}$C spectra of **AzaB-NO**:

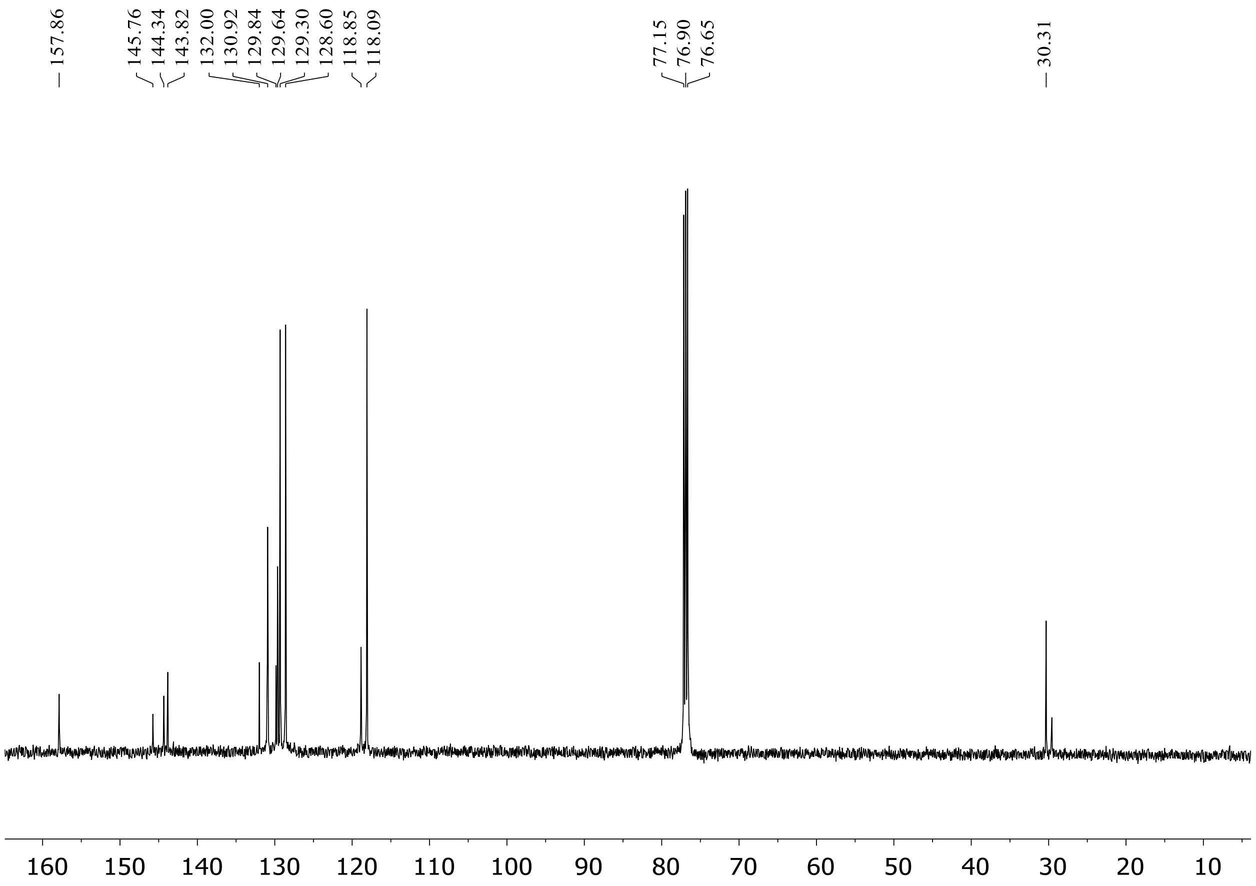

#### HPLC data

High-performance liquid chromatography (HPLC) experiments was performed with Agilent 1260 Infinity II equipped with Poroshell 120 EC-C18 column using gradient elution. The diode array detector was set up for the detection at 670 and 740 nm to obtain clear peaks of the photodonor with and without NO, respectively. First, the transformation of initial form of AzaB-NO to the final form under light was observed, corresponding to the change of the retention time from 14.8 to 14.5 min (on the Figure S*1*, it corresponds to the transformation from black to red spectrum). Then we added the NO donor NONOAte to the reaction mixture and observed backward shift of the peak (marked by blue in the Figure S*1*). It proves the reversibility of the reaction.

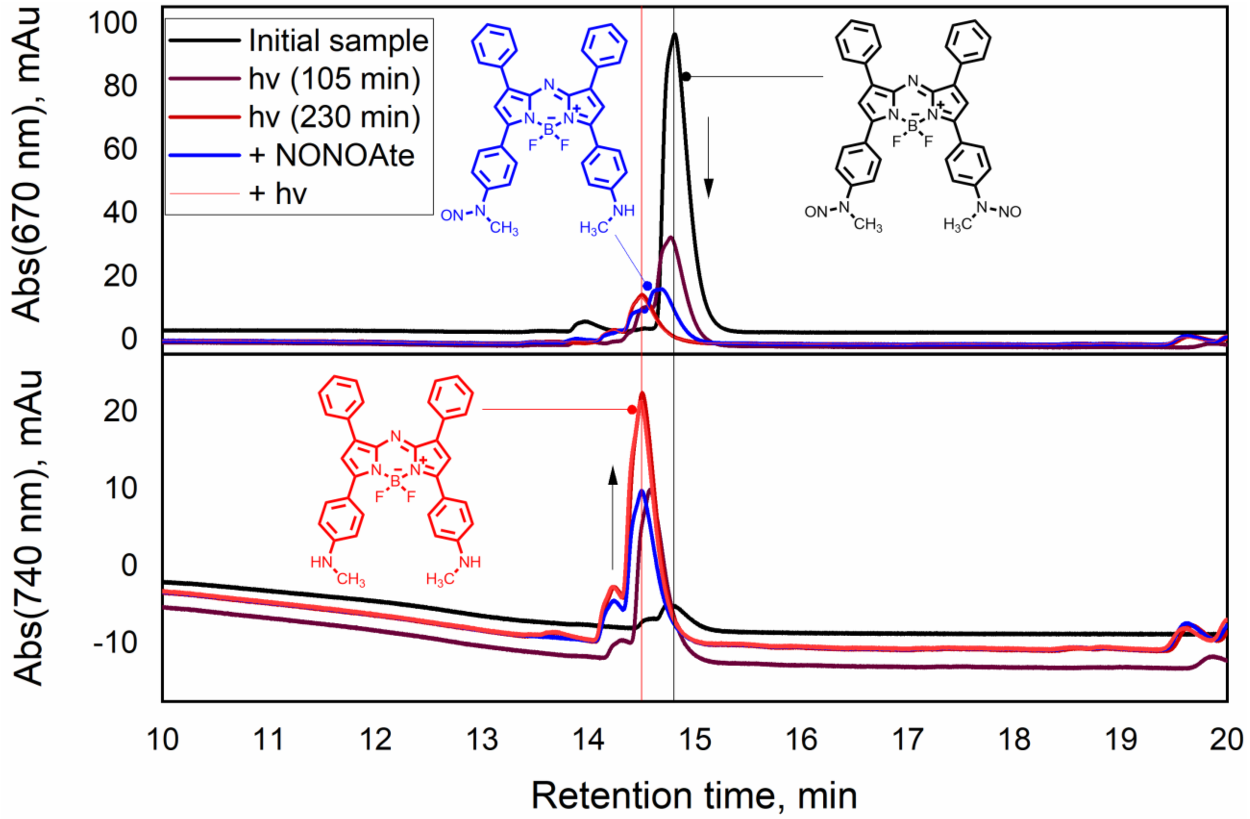

Figure S1. HLPC data for two different channels (670 nm above / 740 nm below). The transformation of initial form of AzaB-NO (absorbing at 670 nm, black spectrum) under light goes to the final form (absorbing at 740 nm, red line). After the addition of NONOate, retention time and absorption of 670/740 nm are shifted to the intermediate state (blue line).

#### Quantification of NO using DAR-2 probe

To measure the yield of NO, we use DAR-2 fluorescent probe. The intensity of fluorescence in any moment is

$$I = I_{\mathrm{exc}}(\phi_0[\mathrm{DAR2}] + \phi_{\mathrm{TZ}}[\mathrm{DAR2_{TZ}}]),$$

where [DAR2] is the concentration of the unaltered probe, $[\mathrm{DAR2_{TZ}}]$ is the concentration of its triazol form (which captured NO), $I_{\mathrm{exc}}$ is the intensity of excitation light, and $\phi_0$ and $\phi_{\mathrm{TZ}}$ are the corresponding proportionality constants. Before the reaction with NO

$$I_0 = I_{\mathrm{exc}}\phi_0[\mathrm{DAR2_0}],$$

where $[\mathrm{DAR2_0}]$ is the initial concentration of DAR-2, and in any moment

$$[\mathrm{DAR2_0}] = [\mathrm{DAR2}] + [\mathrm{DAR2_{TZ}}].$$

Then we have

$$I - I_0 = I_{\mathrm{exc}}(\,\phi_{\mathrm{TZ}} - \phi_0\,)[\mathrm{DAR2_{TZ}}],$$

And finally, dividing by $I_0$, we express the amount of the captured NO as

$$[\mathrm{NO}] = [\mathrm{DAR2_{TZ}}] = [\mathrm{DAR2_0}]\frac{I - I_0}{I_0}\frac{1}{\phi_{\mathrm{TZ}}/\phi_0 - 1}.$$

This equation was used to determine the concentration of NO. The initial concentration of $[\mathrm{DAR2_0}]$ is controlled in experiments, whereas $I$ and $I_0$ are measured fluorescence signals at 550 nm with the excitation at 540 nm. To find the ratio $\phi_{\mathrm{TZ}}/\phi_0$, we measured the fluorescence of DAR-2 in ethanol before and after reaction with the excess of NONOAte and found $\phi_{\mathrm{TZ}}/\phi_0 = 47$, which is close to the ratio of the reported QYs in[1] [1]: $0.34/0.006 \approx 57$ (different buffer was used for measurements).

#### Calibration of the sensor

Calibration of the sensor was done according to the manual, based on the following reaction:

$$2\mathrm{NaNO_2} + 2\mathrm{KI} + 2\mathrm{H_2SO_4} \rightarrow 2\mathrm{NO} + \mathrm{I_2} + 2\mathrm{H_2O} + 2\mathrm{Na_2SO_4}$$

By the addition of certain amount of $NaNO_2$ to the acidic solution we can generate equal amount of NO. Assuming linear response (which is true for smaller concentration, e.g. 100 nM which we used in experiments), we obtain the following ratio: 1 pA corresponds to 5 nM of NO.

[1] Kojima, H.; Hirotani, M.; Nakatsubo, N.; Kikuchi, K.; Urano, Y.; Higuchi, T.; Hirata, Y.; Nagano, T. Bioimaging of Nitric Oxide with Fluorescent Indicators Based on the Rhodamine Chromophore. Anal. Chem. 2001, 73, 1967–1973, doi:10.1021/ac001136i.

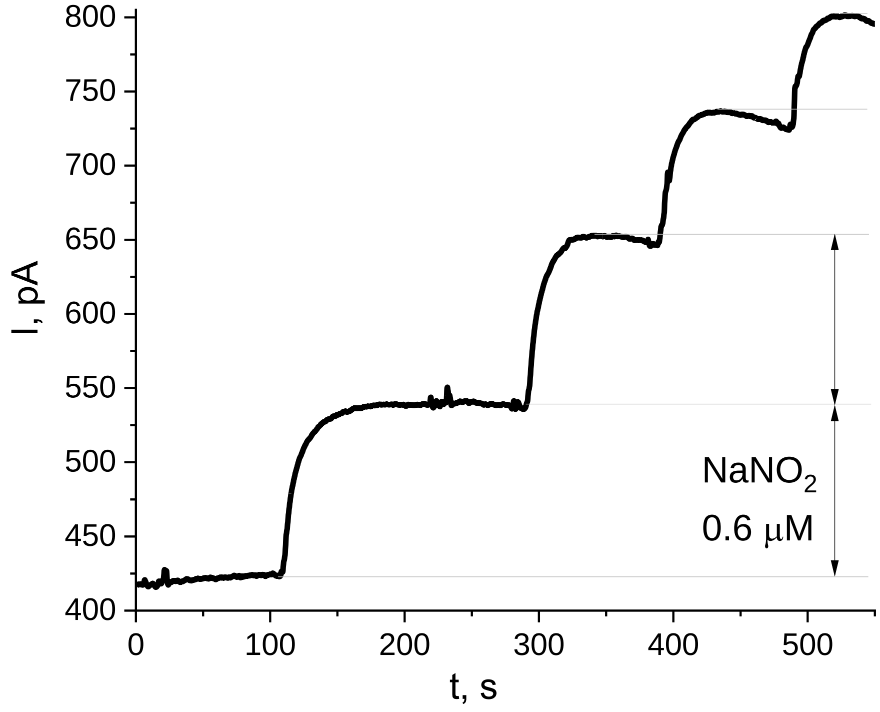

Figure S2. Calibration of the sensor. The response is linear for up to ~1 μM, whereas fast decay of NO is observed at higher concentrations.

#### Evaluation of thermal effects

The changes in temperature due to the absorption of laser radiation was measured experimentally for AzaB-NO photodonor and analogous compound without NO, TP-aza-BODIPY. The results are shown in the figure below.

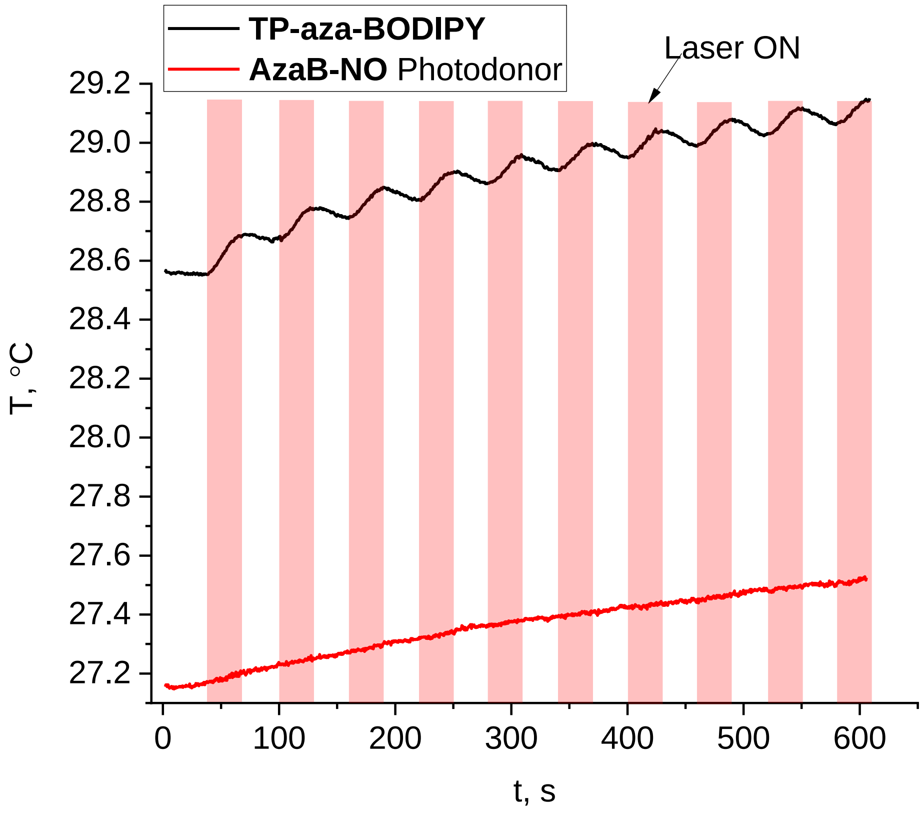

Figure S3. Evaluation of thermal effect. The changes in temperature are comparable for tetraphenyl-aza-BODIPY and **AzaB-NO** photodonor.

#### Model of the NO photorelease

Consider the following kinetic scheme:

$$\begin{cases} A_2 \xrightarrow{k_1} A_0 + 2NO \\ A_0 + NO \xrightarrow{k_2} A_1 \\ A_1 + NO \xrightarrow{k_3} A_2 \end{cases}$$

where $A_2$ is the concentration of photodonor, $A_1$ is the concentration of dye with only one N-Nitroso bond left, and $A_0$ is the concentration of dye not bearing any NO. $NO$ denotes the concentration of NO, the forward rate constant $k_1$ depends on light intensity and the reverse rate constants $k_2, k_3$ are intrinsic properties of the molecule. Note that we assume that two NO molecules dissociate at once for simplicity. For now, we do not consider the degradation of NO assuming it much slower. We obtain the following:

$$\begin{cases} \dfrac{dA_2}{dt} = -k_1 \cdot A_2 + k_3 \cdot A_1 \cdot NO \\ \dfrac{dA_1}{dt} = k_2 \cdot A_0 \cdot NO - k_3 \cdot A_1 \cdot NO \\ \dfrac{dA_0}{dt} = k_1 \cdot A_2 - k_2 \cdot A_0 \cdot NO \\ \dfrac{dNO}{dt} = 2k_1 \cdot A_2 - k_2 \cdot A_0 \cdot NO - k_3 \cdot A_1 \cdot NO \end{cases}$$

There is also the following condition relating free NO with $A_0$ and $A_1$:

$$NO = A_1 + 2A_0$$

Now consider the stationary case (time derivatives = 0), then the system of differential equations together with the above equation give:

$$\begin{cases} A_0 = NO\left(2 + \dfrac{k_2}{k_3}\right)^{-1} \\ A_1 = \dfrac{k_2}{k_3} A_0 = \dfrac{k_2}{k_3} NO\left(2 + \dfrac{k_2}{k_3}\right)^{-1} \\ A_2 = \dfrac{k_2}{k_1} NO \cdot A_0 = \dfrac{k_2}{k_1} NO^2\left(2 + \dfrac{k_2}{k_3}\right)^{-1} \end{cases},$$

There is also the initial conditions:

$$[A_2]_0 = A_2 + A_1 + A_0,$$

where $[A_2]_0$ is $A_2$ before the reaction (i.e. before the photodecomposition begins). Substituting the above expressions, we have the following quadratic equation for NO:

$$\frac{k_2}{k_1}NO^2 + NO\left(1+\frac{k_2}{k_3}\right) - [A_2]_0\left(2+\frac{k_2}{k_3}\right)$$

This equation gives the following solution

$$NO = \frac{k_1}{2k_2}\left(-\left(1+\frac{k_2}{k_3}\right) + \sqrt{\left(1+\frac{k_2}{k_3}\right)^2 + 4\frac{k_2}{k_1}\left(2+\frac{k_2}{k_3}\right)[A_2]_0}\right)$$

Let's discuss the limiting cases. Letting $k_1$ tend to zero, we get the case when there is no direct reaction. Accordingly, NO is not generated and the concentration tends to zero as well. The rate constant $k_1$ is obviously proportional to the local light intensity (this result can be, for instance, recovered from the Beer-Lambert Law decomposed for infinitely small thickness of absorbing layer).

To consider the case $k_2 \to 0$, it is necessary to use the Taylor series expansion with a small parameter $\frac{k_1}{2k_2}$:

$$NO \approx \frac{k_1}{2k_2}\left(1+\frac{k_2}{k_3}\right)\left(-1+1+2\frac{k_2}{k_1}\frac{\left(2+\frac{k_2}{k_3}\right)}{\left(1+\frac{k_2}{k_3}\right)^2}[A_2]_0\right) = \frac{\left(2+\frac{k_2}{k_3}\right)}{\left(1+\frac{k_2}{k_3}\right)}[A_2]_0.$$

Further behavior depends on $\frac{k_2}{k_3}$, but obviously the maximal results $2[A_2]_0$ is achieved at $k_2 = 0$. So, in the absence of a reverse reaction, the concentration of NO tends to the doubled initial concentration of the donor.

#### Hardware

Output of WPI TBR-1025 free radical analyzer is the analogous voltage. An ADS1115 16-bit ADC module was used to convert it into the digital value. The ADC module was connected to Arduino Nano board using standard jumper wires and solderless breadboard. The overall hardware connections are listed in the following figure.

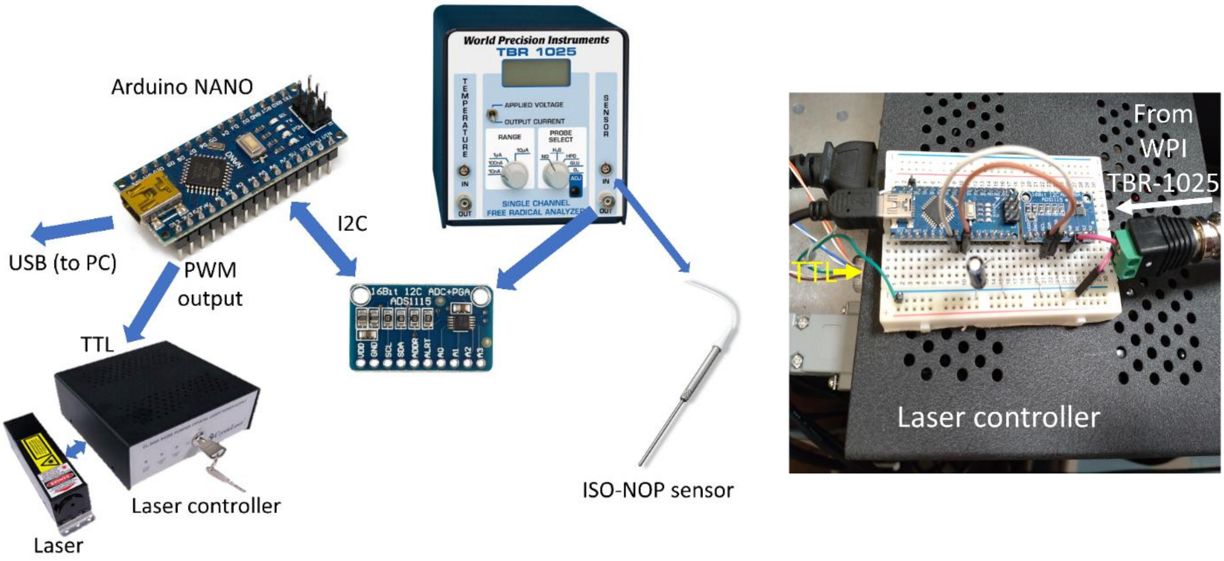

Figure S4. Overview of the system's hardware.

Arduino microcontroller was set up to establish the serial-port connection with PC via the USB and to send the readings upon request (`m` – NO sensor, `t` – temperature sensor). Special commands were reserved to set up gain (the ADS1115 module has a built-in amplifier with variable gain). Additionally, one of pulse-width modulation (PWM) pins of Arduino microcontroller was connected to the TTL input of the laser controller. The output of this pin was controlled by special command arriving from PC. The firmware for the microcontroller is listed below.

```
#include <Adafruit_ADS1X15.h>
int inByte = 0;
int gain = 0;
Adafruit_ADS1X15 ADC1;
int level = 0;
int LEDpin = 11;
void setup() {
  digitalWriteFast(LEDpin,LOW);
  Serial.begin(9600);
  if(!ADC1.begin(0x48)) Serial.println("Cannot start ADS1115");
  ADC1.setGain(GAIN_TWOTHIRDS);
}
void loop() {
  if (Serial.available() > 0) {
    inByte = Serial.read();
    if(inByte == 'm') {
      Serial.println(ADC1.readADC_SingleEnded(0));
    }
    if (inByte == 't'){
      Serial.println(ADC1.readADC_SingleEnded(2));
    }
    if(inByte == 'u') {
       level = Serial.parseInt();
       analogWrite(LEDpin,level);
    }
    if(inByte == 'd') {
       analogWrite(LEDpin,LOW);
    }
    if(inByte == 'g') {
      gain = Serial.parseInt();
```

```
        switch(gain) {
          case 1:
            ADC1.setGain(GAIN_ONE);
            break;
          case 2:
            ADC1.setGain(GAIN_TWO);
            break;
          case 4:
            ADC1.setGain(GAIN_FOUR);
            break;
          case 8:
            ADC1.setGain(GAIN_EIGHT);
            break;
          case 16:
            ADC1.setGain(GAIN_SIXTEEN);
            break;
          default:
            ADC1.setGain(GAIN_TWOTHIRDS);
            break;
        }
      }
    }
  }
```
